# Mechanical Properties of White Matter Tracts in Aging Assessed via Anisotropic MR Elastography

**DOI:** 10.1101/2024.05.08.593260

**Authors:** Diego A. Caban-Rivera, Lance T. Williams, Matthew D. J. McGarry, Daniel R. Smith, Elijah E. W. Van Houten, Keith D. Paulsen, Philip V. Bayly, Curtis L. Johnson

## Abstract

Magnetic resonance elastography (MRE) is a promising neuroimaging technique to probe tissue microstructure, which has revealed widespread softening with loss of structural integrity in the aging brain. Traditional MRE approaches assume mechanical isotropy. However, white matter is known to be anisotropic from aligned, myelinated axonal bundles, which can lead to uncertainty in mechanical property estimates in these areas when using isotropic MRE. Recent advances in anisotropic MRE now allow for estimation of shear and tensile anisotropy, along with substrate shear modulus, in white matter tracts. The objective of this study was to investigate age-related differences in anisotropic mechanical properties in human brain white matter tracts for the first time. Anisotropic mechanical properties in all tracts were found to be significantly lower in older adults compared to young adults, with average property differences ranging between 0.028-0.107 for shear anisotropy and between 0.139-0.347 for tensile anisotropy. Stiffness perpendicular to the axonal fiber direction was also significantly lower in older age, but only in certain tracts. When compared with fractional anisotropy measures from diffusion tensor imaging, we found that anisotropic MRE measures provided additional, complementary information in describing differences between the white matter integrity of young and older populations. Anisotropic MRE provides a new tool for studying white matter structural integrity in aging and neurodegeneration.

## 1. INTRODUCTION

Magnetic resonance elastography (MRE) is a noninvasive MR imaging technique that produces quantitative maps of viscoelastic mechanical properties in soft biological tissues (Hiscox et al., 2016; Muthupillai et al., 1995; Sack, 2022). MRE experiments are performed with an external vibration source used to propagate shear waves through the skull into the brain, inducing micron-level displacements. The MRI-captured wave motion images are then inverted to generate maps of mechanical properties of the tissue (Manduca et al., 2021). In the human brain, MRE has been utilized in several studies revealing softening of the neural tissue in aging and in neurological diseases, including Alzheimer’s disease and other dementias (Delgorio et al., 2023; Hiscox et al., 2020; Huston et al., 2016; Murphy et al., 2019; Pavuluri et al., 2023). Through preclinical animal models, changes to mechanical properties of brain tissue have been shown to relate to the composition and organization of microscale elements, indicated by significant correlations with neuronal density, myelin content and oligodendrocytes, microglia, and astrocytes (Freimann et al., 2013; Schregel et al., 2012; Weickenmeier et al., 2016, 2017). MRE measures of stiffness and viscosity can assess these microscale changes in tissue composition and organization that are affected by disease (Sack et al., 2013).

Effects of aging on brain tissue integrity have been studied with MRE, with several reports of tissue softening in older age (Hiscox et al., 2021). Most investigations have focused on softening globally or in large regions (Arani et al., 2015; Sack et al., 2009, 2011; Takamura et al., 2020), or local effects in smaller gray matter regions (Delgorio et al., 2021; Hiscox et al., 2018), with an average reported annual decrease in stiffness of 0.8% (or 0.006-0.025 kPa/yr), and with significant variability in apparent age-related rate of softening depending on the region. However, MRE studies explicitly considering effects of age or age-related neurodegenerative conditions on white matter (WM) mechanical properties have been limited, likely due, in part, to the need for an anisotropic MRE technique to resolve direction-dependent mechanical properties in WM. Directional differences in properties arise from aligned axon fibers, and can lead to errors and uncertainty in traditional, isotropic MRE inversions (Anderson et al., 2016; McGarry et al., 2021).

Early attempts to characterize mechanical properties in WM tracts assumed mechanical isotropy during inversion (Guo et al., 2013; Johnson et al., 2013), but anisotropic inversion methods have recently been developed to better capture the directionally-dependent mechanical behavior of WM (McGarry et al., 2021; Romano et al., 2012). These anisotropic methods have varied with regards to the underlying material model, the number of parameters estimated, and the numerical inversion schemes, with each of these elements impacting the overall performance of the MRE reconstruction. An early two-parameter approach that models shear modulus parallel and perpendicular to the fiber direction was initially applied to study breast (Sinkus et al., 2005) and skeletal muscle (Green et al., 2013); this approach does not capture the potential tensile anisotropy of brain tissue. Conversely, an orthotropic, nine-parameter model was applied to WM and the corticospinal tract specifically (Romano et al., 2012, 2014); due to the large number of coefficients of dramatically different magnitudes, estimating these parameters accurately is difficult from traditional MRE data (Miller et al., 2018a, 2018b). Recently, a three-parameter, nearly-incompressible, transversely-isotropic material model was proposed that comprises a substrate shear modulus and two anisotropy parameters that represent the differences in Young’s modulus and shear modulus relative to the assumed fiber direction. This model has been shown to describe WM mechanics with a minimal number of parameters (Feng et al., 2013). We have integrated this model into the finite element-based, nonlinear inversion algorithm (NLI) (McGarry et al., 2012) – termed transversely-isotropic NLI (TI-NLI) (McGarry et al., 2021, 2022) – to estimate WM anisotropic properties while also accounting for heterogeneity in properties and varying fiber direction throughout the brain.

To improve the accuracy and stability of property estimates with TI-NLI, we use multi-excitation MRE to produce diverse displacement data (Anderson et al., 2016; Smith et al., 2020). Shear waves in fibrous, transversely-isotropic materials travel at different wave speeds based on wave propagation and polarization directions (Tweten et al., 2015), and, as such, MRE displacement data must deform the tissue of interest in multiple directions in order to estimate anisotropic parameters reliably (Tweten et al., 2017). Multi-excitation MRE uses two or more vibration sources, applied sequentially, to generate different wave patterns throughout the brain (Smith et al., 2020), and, when coupled with TI-NLI, produces repeatable anisotropic property estimates in WM tracts (Smith et al., 2022).

In this study, we estimate anisotropic mechanical properties of WM tracts of a younger and older population to evaluate how these parameters are affected by aging. MRE outcomes of substrate shear stiffness, shear anisotropy, and tensile anisotropy are compared between groups of older adults and younger adults. We also investigate how age-related differences in mechanical properties from anisotropic MRE compare with metrics from diffusion MRI, which is commonly used to assess WM integrity, and has shown lower fractional anisotropy in aging (Davis et al., 2009; Lebel et al., 2012; Michielse et al., 2010).

## 2. METHODS

### 2.1 Participant information

Participants included 20 younger adults (25.2 ± 2.1 years, 11 females) and 19 older adults (68.1 ± 5.9 years, 8 females). Each participant provided informed, written consent to participate in this study approved by the Delaware Institutional Review Board. All participants were screened prior to enrollment to exclude those with any history of neurological disease or brain injury. One older participant did not complete the entire scan session successfully and this data set was excluded from the study (final older adult group, N=18). Each participant completed an MR imaging protocol on a Siemens 3T Prisma MRI scanner with a 20-channel head coil.

### 2.2 Image Acquisition and Processing

#### Multi-excitation MR Elastography

Mechanical shear waves were introduced into the brain at 50 Hz using a pneumatic system (Resoundant, Inc.; Rochester, MN) with two vibration sources: a pillow driver that applies anterior-posterior (AP) motion and a custom designed left-right (LR) actuator highlighted in Figure 1A, applied separately, resulting in two distinct, three-dimensional wave fields for inversion (Figure 1C) (Smith et al., 2020). The LR actuator combines a silicone bottle driver with 3D printed components that affix the actuator to the head coil to enable stable vibrations and optimal coupling against the participants’ temple (Caban-Rivera et al., 2022; Kailash et al., 2019). Wave fields from both actuations were captured with a 3D multiband, multishot spiral sequence (Johnson et al., 2016; McIlvain et al., 2022) that encodes the full vector displacement fields generated in the brain by external actuation. Imaging parameters for each acquisition included 4 phase offsets, 2.0 mm isotropic voxel resolution, 240 x 240 mm^2^ FOV, 64 slices, TR/TE = 2240/76 ms. Shear wave displacement fields were calculated via subtraction of opposite-polarity MRE phase images, phase unwrapping was performed using FSL PRELUDE (Jenkinson, 2003), and temporal Fourier filtering was applied to isolate 50 Hz harmonic motion.

**Figure 1:**
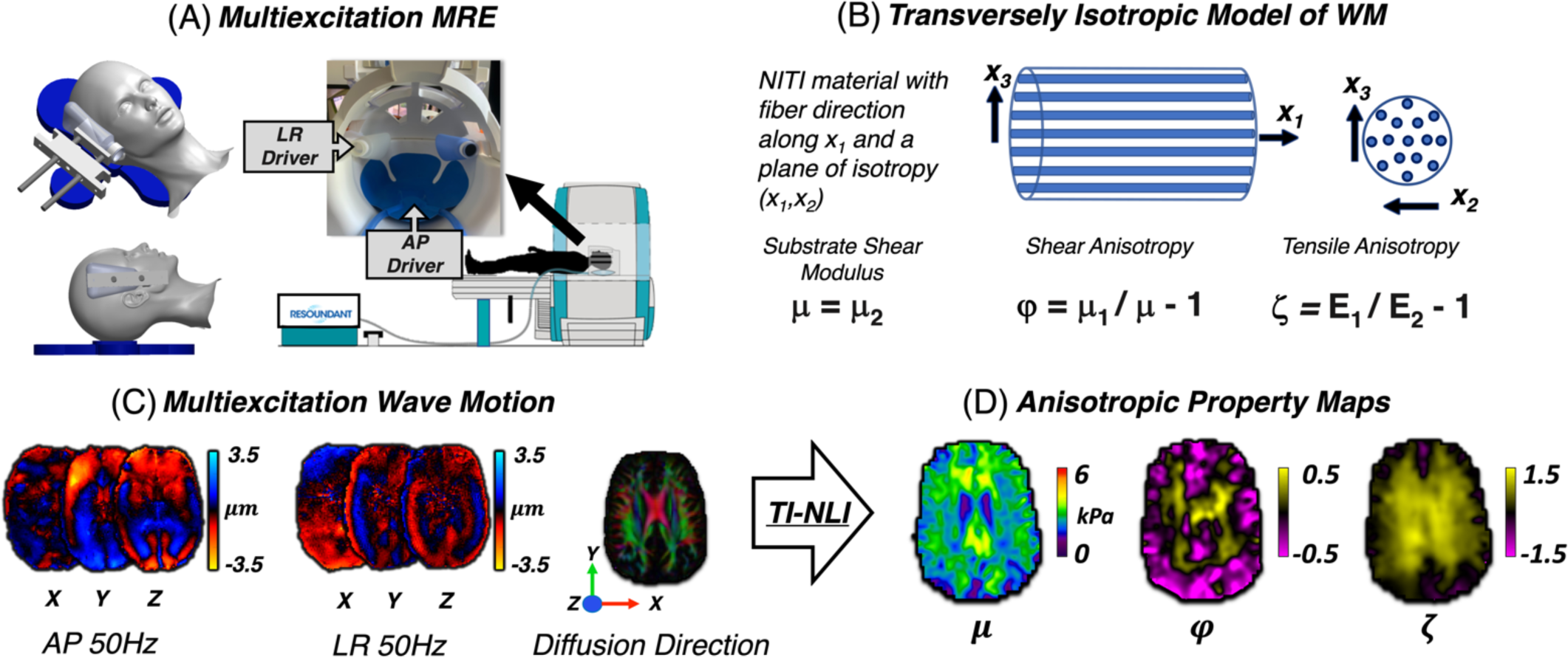
Overview of multi-excitation MRE and TI-NLI. A) Depiction of the experimental set up showing the custom designed left-right (LR) bottle actuator, and the anterior-posterior (AP) pillow driver. B) Schematic of a transversely isotropic material comprising aligned fibers in a substrate, representing fibrous white matter. C) Inputs to for TI-NLI: AP and LR wave fields at 50 Hz and fiber directions from diffusion MRI. D) Example anisotropic parameter maps reconstructed with TI-NLI.

#### Anatomical T1-Weighted Images

High resolution T1-weighted anatomical images were acquired with an MPRAGE sequence for registration and segmentation with the following imaging parameters: 0.9 mm isotropic voxels; 256 x 256 mm^2^ FOV; 176 slices; repetition time (TR)/echo time (TE)/inversion time (TI) = 2300/2.32/900 ms. T1-weighted images were processed using the *fsl_anat* script from FSL (Jenkinson et al., 2012) to automatically segment and register structural images to standard space templates. Images were processed with the following steps: 1) reorientation to MNI standard orientation, 2) bias-field correction with FSL FAST, 3) linear and nonlinear registration to standard space (FLIRT and FNIRT), 4) binary mask created via FNIRT-based brain-extraction, and 5) tissue type segmentation. Bias-corrected, brain-extracted T1-weighted images outputted from *fsl_anat* were registered to MRE images, and the binary mask was applied to the MRE magnitude images prior to calculating the shear wave displacements and estimating mechanical properties.

#### Diffusion MRI

Diffusion-weighted images were acquired with a simultaneous, multi-slice EPI sequence (210 x 240 x 138 mm^3^ field-of-view, 1.5 mm isotropic resolution, 92 slices, TR/TE = 3520/95.2 ms, b-values = 1500, 3000 s/mm^2^ over 128 directions). Reference scans were acquired with opposite phase encoding direction and zero b-value for distortion corrections using TOPUP from FSL (Andersson et al., 2003; Smith et al., 2004). Diffusion images were registered linearly to MRE magnitude images with FSL FLIRT (Jenkinson et al., 2002; Jenkinson & Smith, 2001), and the diffusion gradient directions for each image were rotated accordingly to correct for motion between scans. Diffusion tensors in each voxel were then estimated using FSL Diffusion Toolbox (FDT) which outputs eigenvectors and eigenvalues of the diffusion tensor and associated maps of fractional anisotropy (FA). The first principal eigenvector of the diffusion tensor (the direction of maximal diffusivity) is interpreted as the local fiber axis and used in the inversion algorithm for estimating anisotropic parameters.

### 2.3 Transversely Isotropic Nonlinear Inversion (TI-NLI)

Two distinct displacement fields from multi-excitation MRE, including shear waves with distinct propagation and polarization directions, and the primary eigenvector from diffusion MRI were input to TI-NLI to estimate a single set of anisotropic mechanical parameters (McGarry et al., 2021, 2022). Briefly, NLI is a finite-element based optimization composed of two discrete problems: first the forward problem calculates the expected displacement fields from an estimated set of material properties, followed by the inverse problem that iteratively updates the unknown material property distribution based on the difference between measured and calculated displacements (McGarry et al., 2012; Van Houten et al., 1999). TI-NLI incorporates a finite element implementation of a heterogeneous, nearly incompressible, transversely isotropic model to describe behavior of brain tissue (Feng et al., 2013; Tweten et al., 2015). Outputs from TI-NLI include spatial maps of three model parameters that describe the behavior of viscoelastic, fiber-reinforced, anisotropic tissue (Figure 1D):

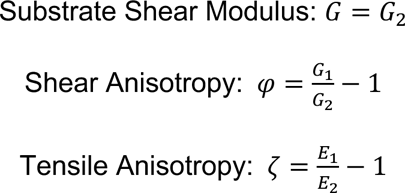

Here, *G* refers to the complex valued substrate shear modulus and Young’s modulus is denoted by *E*, while subscripts 1 and 2 refer to components parallel and perpendicular to the fiber axis, respectively. Additionally, we compute substrate shear stiffness as a composite parameter of the substrate shear modulus 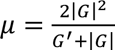 which is proportional to the square of the wave speed in a viscoelastic material and is a measure commonly reported in MRE literature.

### 2.4 Tract-Based Spatial Statistics

We investigated age-based differences in WM mechanical properties through a voxel-wise analysis using tract-based spatial statistics (TBSS) (Smith et al., 2006). TBSS accounts for alignment inconsistencies across subjects that can confound voxel-wise statistics by generating a common WM skeleton for voxel-wise analysis between subjects. FA maps from diffusion MRI underwent preprocessing including erosion, removal of outlier slices, and nonlinear registration to an average FA template (*FMRIB58_FA*) in standard space at 1 mm isotropic resolution, with voxels in which FA > 0.2 were included. Reconstructed MRE property maps (μ, ϕ, ζ), in the same native space as FA, were then projected onto the mean FA skeleton using *tbss_non_fa* in FSL for statistical analyses in standard space. Voxel-wise statistical analyses were performed with FSL *randomize* to test for differences between younger and older adults for all mechanical parameters, with significance level α = 0.05. We calculated the percentage of voxels in the skeleton that exhibited significant differences between groups. We also compared the statistical maps for each of the MRE measures and FA from diffusion, calculating overlap in significant voxels describing differences between age groups.

### 2.5 ROI-Based Analysis of WM Tracts

We performed region-of-interest (ROI) analyses using WM tract masks to confirm the TBSS findings. Tract masks were obtained from WM atlases (Hua et al., 2008; Mori et al., 2008) and applied to the TBSS outputs in standard space. Tracts of interest included: corpus callosum body (CCB), forceps major (Fmaj), forceps minor (Fmin), corona radiata (CR), corticospinal tract (CST), anterior thalamic radiation (ATR), posterior thalamic radiation (PTR), and superior longitudinal fasciculus (SLF). Average property values were calculated within each tract mask for voxels that coincide with the WM skeleton from TBSS. Two-sample t-tests compared properties between younger and older adults for each tract with Bonferroni correction for multiple comparisons. We additionally employed a logistic regression analysis to determine the novel information captured by anisotropic MRE relative to diffusion MRI parameters. FA from diffusion for a given tract was added in the model as a classifier first, followed by either ϕ or ζ added second. A significant effect of adding MRE in classifying age group indicates that the mechanical anisotropy adds additional information beyond what is captured by diffusion alone.

## 3. RESULTS

Representative property maps for one younger adult (30 years, female) and one older adult (61 years, female) are shown in Figure 2 for qualitative comparison of the anisotropic mechanical properties and their distribution in the whole brain. In general, the younger adult shows higher whole brain substrate stiffness, while shear anisotropy and tensile anisotropy show similar structure between ages, with the older adult exhibiting lower anisotropy in the WM regions. Table 1 provides a summary of the findings for the anisotropic MRE parameters with mean and standard deviations for each WM tract.

**Figure 2:**
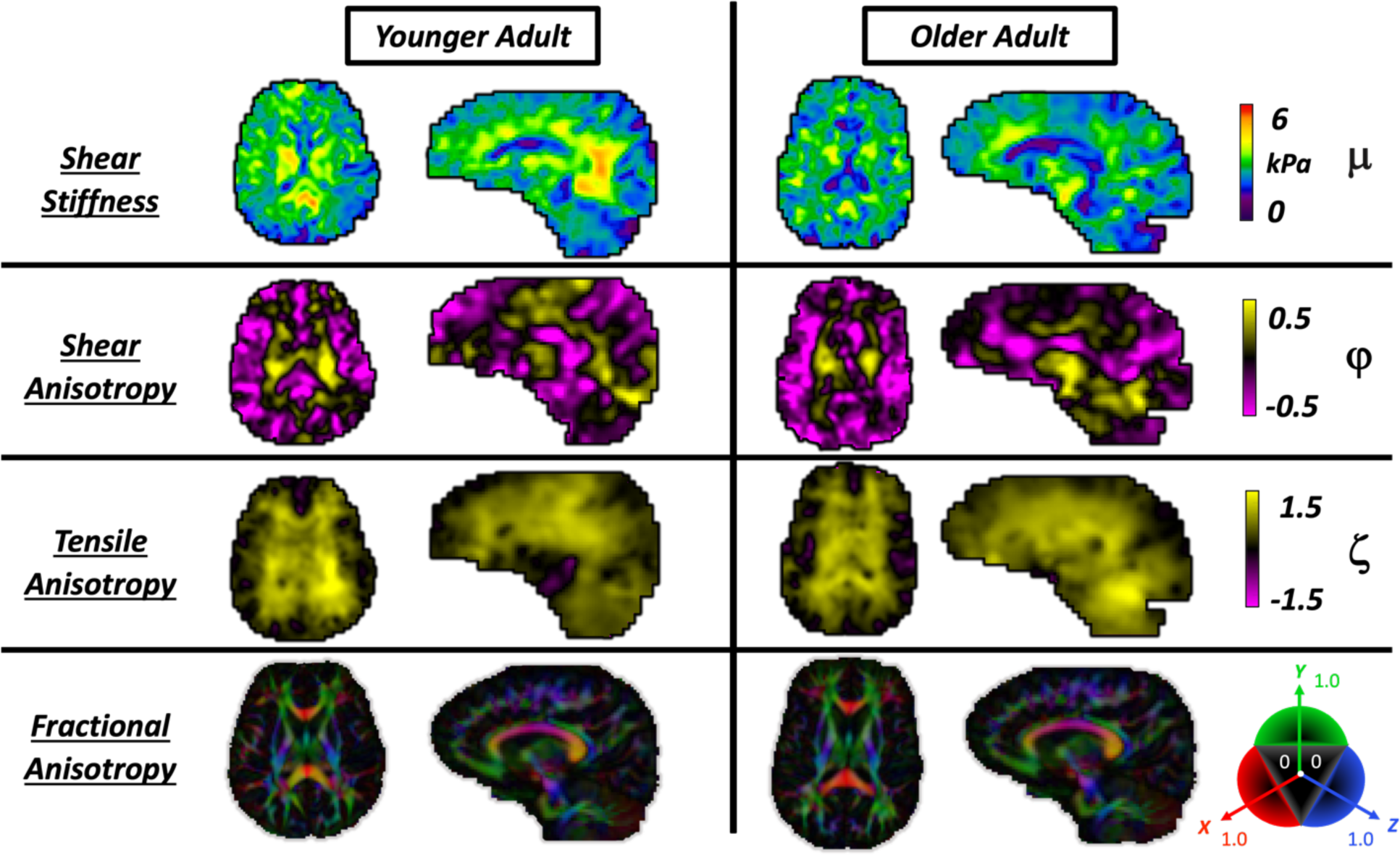
Comparison of representative whole brain property maps between one younger adult (30 years, F) and one older adult (61 years, F). Rows 1-3 illustrate outputs from the transversely isotropic nonlinear inversion, where anisotropic parameters have similar structural contrast between the groups, but lower values across the whole brain in the older adults. The 4th row shows the colored fractional anisotropy map estimated from the diffusion tensor with the corresponding direction-color combinations: x-red, y-green, and z-blue. We observed lower values for global stiffness, anisotropy, and FA in the older adults.

**Table 1:** Overview of TI-NLI property means and standard deviations in white matter tracts for young adults (YA) versus older adults (OA). Statistically significant group differences after Bonferroni correction are bolded and starred for significance level **p < 0.00625**.

| Region | Shear Stiffness (kPa) |  |  | Shear Anisotropy |  |  | Tensile Anisotropy |  |  |
| --- | --- | --- | --- | --- | --- | --- | --- | --- | --- |
|  | YA | OA |  | YA | OA |  | YA | OA |  |
| **CCB** | 2.45 ± 0.29 | 2.44 ± 0.35 | $p = 0.890$ | 0.26 ± 0.10 | 0.19 ± 0.10 | $p = 0.041$ | 1.10 ± 0.16 | 0.83 ± 0.13 | **$p < 0.001^*$** |
| **Fmaj** | 2.72 ± 0.19 | 2.62 ± 0.19 | $p = 0.040$ | 0.00 ± 0.09 | -0.07 ± 0.12 | $p = 0.038$ | 1.11 ± 0.19 | 0.97 ± 0.13 | $p = 0.013$ |
| **Fmin** | 2.95 ± 0.22 | 2.62 ± 0.24 | **$p < 0.001^*$** | 0.05 ± 0.05 | -0.05 ± 0.08 | **$p < 0.001^*$** | 0.88 ± 0.20 | 0.53 ± 0.19 | **$p < 0.001^*$** |
| **CR** | 2.69 ± 0.13 | 2.61 ± 0.18 | $p = 0.110$ | 0.21 ± 0.03 | 0.12 ± 0.05 | **$p < 0.001^*$** | 1.06 ± 0.13 | 0.78 ± 0.12 | **$p < 0.001^*$** |
| **CST** | 2.82 ± 0.21 | 2.77 ± 0.23 | $p = 0.537$ | 0.35 ± 0.07 | 0.25 ± 0.06 | **$p < 0.001^*$** | 1.12 ± 0.13 | 0.88 ± 0.14 | **$p < 0.001^*$** |
| **ATR** | 3.00 ± 0.16 | 2.78 ± 0.21 | **$p < 0.001^*$** | 0.09 ± 0.03 | 0.05 ± 0.04 | **$p < 0.001^*$** | 1.02 ± 0.16 | 0.77 ± 0.13 | **$p < 0.001^*$** |
| **PTR** | 2.87 ± 0.18 | 2.74 ± 0.20 | $p = 0.032$ | -0.05 ± 0.10 | -0.08 ± 0.07 | $p = 0.336$ | 1.18 ± 0.19 | 1.02 ± 0.08 | **$p = 0.003^*$** |
| **SLF** | 2.71 ± 0.13 | 2.67 ± 0.16 | $p = 0.294$ | 0.10 ± 0.03 | 0.03 ± 0.04 | **$p < 0.001^*$** | 1.03 ± 0.14 | 0.80 ± 0.09 | **$p < 0.001^*$** |

### 3.1 Substrate Shear Stiffness (μ)

Figure 3 presents differences in substrate shear stiffness between groups. Through voxel-wise analysis with TBSS, we observed various middle to anterior regions where stiffness was significantly lower in older adults compared to younger adults (6.1% of the WM skeleton). Through ROI analysis, we observed that the older adults group exhibited significantly lower stiffness in the Fmin (2.61 vs. 2.95 kPa; p < 0.001) and in the ATR (2.78 vs. 3.00 kPa; p < 0.001). No voxels from TBSS or ROIs appeared significantly stiffer in older adults compared to younger adults.

**Figure 3:**
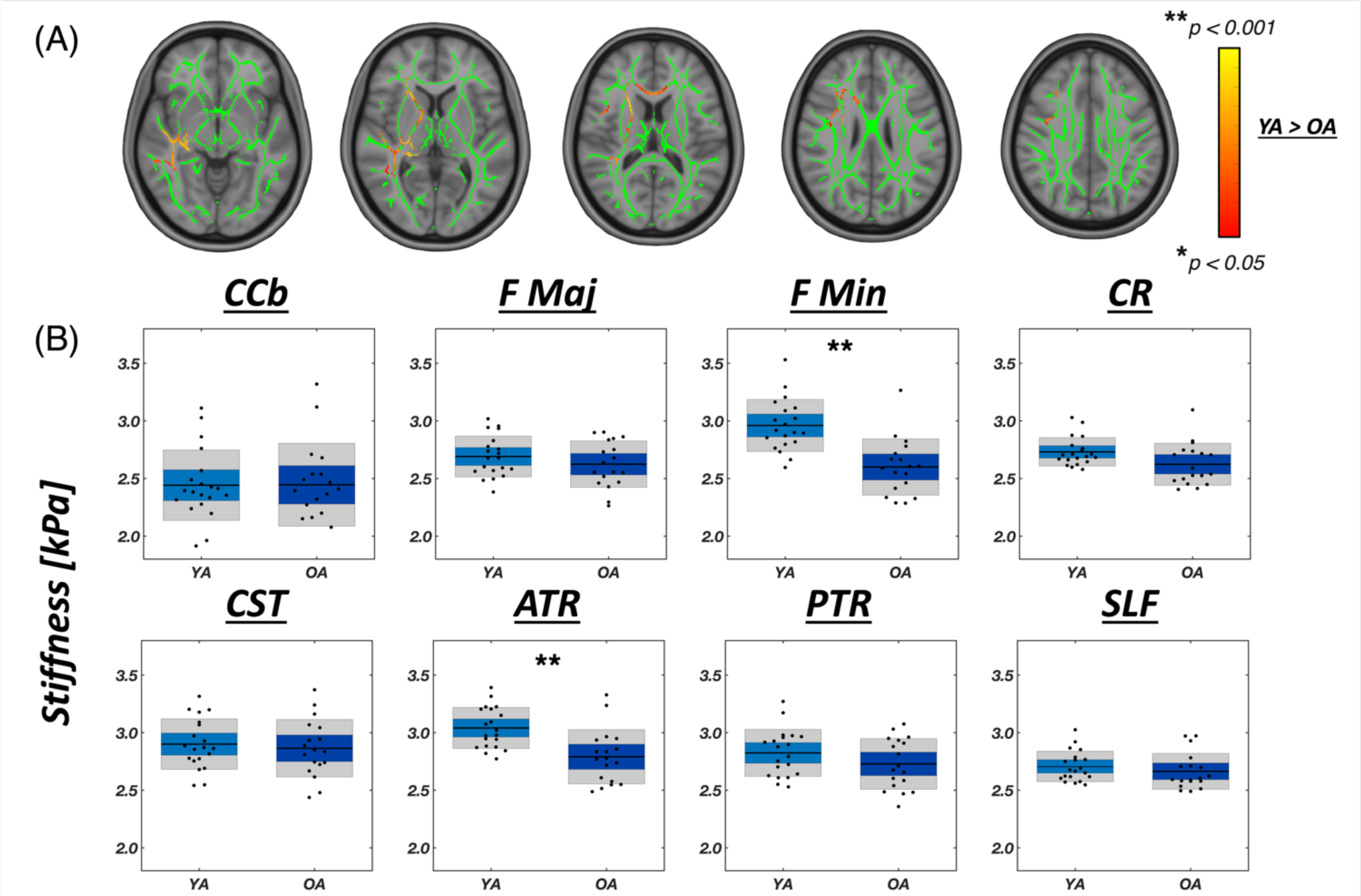
(A) Results from voxel-wise statistical analysis comparing younger adults (YA) versus older adults (OA) for substrate shear stiffness (μ), depicted for several axial slices (Z = -6, 5, 14, 26, 35). Significant voxels from the YA>OA contrast are highlighted in red-yellow, overlayed on the white matter skeleton shown in green. (B) Segmented white matter region averages of shear stiffness, with asterisks indicating level of statistical significance for two-sample student t-tests with Bonferroni correction, * p<0.00625, ** p<0.001.

### 3.2 Shear Anisotropy (ϕ)

Figure 4 presents differences in shear anisotropy between groups. TBSS analysis showed many voxels in WM, particularly in the corpus callosum and periventricular regions, exhibited lower shear anisotropy in older adults (41.8%). Lower shear anisotropy indicates that the ratio of shear stiffness parallel to fiber direction relative to the stiffness perpendicular to fibers is smaller. Since most tracts exhibit a positive ϕ in tracts in younger adults, lower anisotropy in older adults indicates that the stiffness parallel to fibers is more similar to the perpendicular stiffness with age. All tract ROIs showed significantly lower ϕ in older adults except the CCB (0.19 vs. 0.26; p = 0.041), Fmaj (-0.07 vs 0.00; p = 0.038), and PTR (-0.08 vs. -0.05; p = 0.336). The highest shear anisotropy was seen in the CST for both groups – 0.35±0.07 in younger adults and 0.25±0.06 in older adults – which also showed the greatest difference between groups (p < 0.001). No voxels from TBSS or ROIs appeared to have significantly higher shear anisotropy in older adults compared to younger adults.

**Figure 4:**
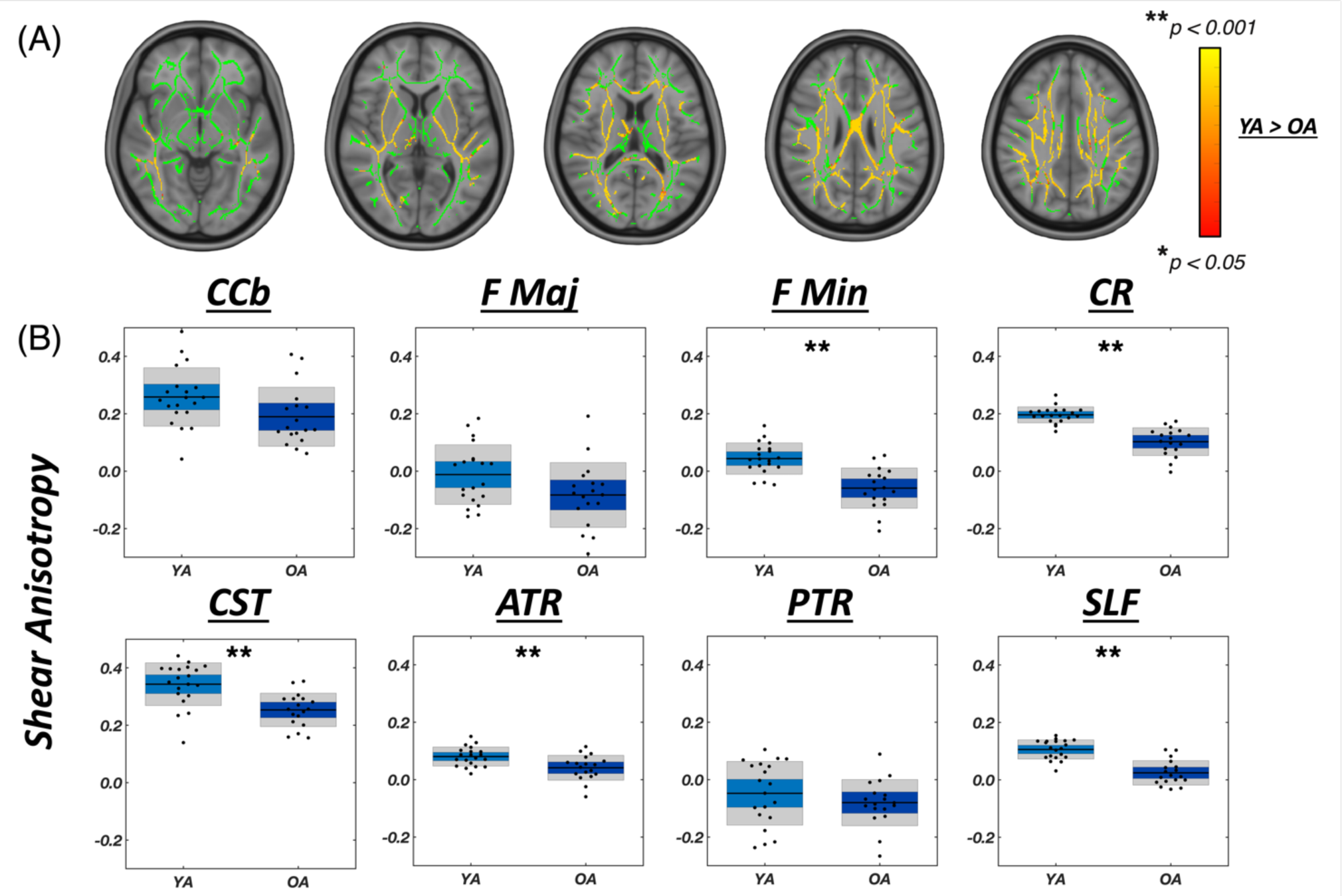
(A) Results from voxel-wise statistical analysis comparing younger adults (YA) versus older adults (OA) for substrate shear stiffness (ϕ), depicted for several axial slices (Z = -6, 5, 14, 26, 35). Significant voxels from the YA>OA contrast are highlighted in red-yellow, overlayed on the white matter skeleton shown in green. (B) Segmented white matter region averages of shear anisotropy, with asterisks indicating level of statistical significance for two-sample student t-tests with Bonferroni correction, * p<0.00625, ** p<0.001.

### 3.3 Tensile Anisotropy (ζ)

Figure 5 presents differences in tensile anisotropy between groups. TBSS analysis showed that most voxels of the cerebrum exhibited lower ζ in older adults (59.8% of total skeleton). Older adults exhibited lower ζ in all WM tract ROIs investigated (all p < 0.00625) except the Fmaj (0.97 vs. 1.11; p = 0.013), with significant differences in anisotropy between young and old ranging from -0.16 (PTR: -14.3%; p = 0.003) to -0.35 (Fmin: -49.0%; p < 0.001). Tensile anisotropy in WM tracts in younger adults is near 1, indicating that the tensile modulus is approximately double in the fiber direction (Smith et al., 2022); lower ζ indicates tensile modulus in the fiber direction is lower in older adults. As with stiffness and shear anisotropy, no voxels from TBSS or ROIs appeared to have significantly higher tensile anisotropy in older adults compared to younger adults.

**Figure 5:**
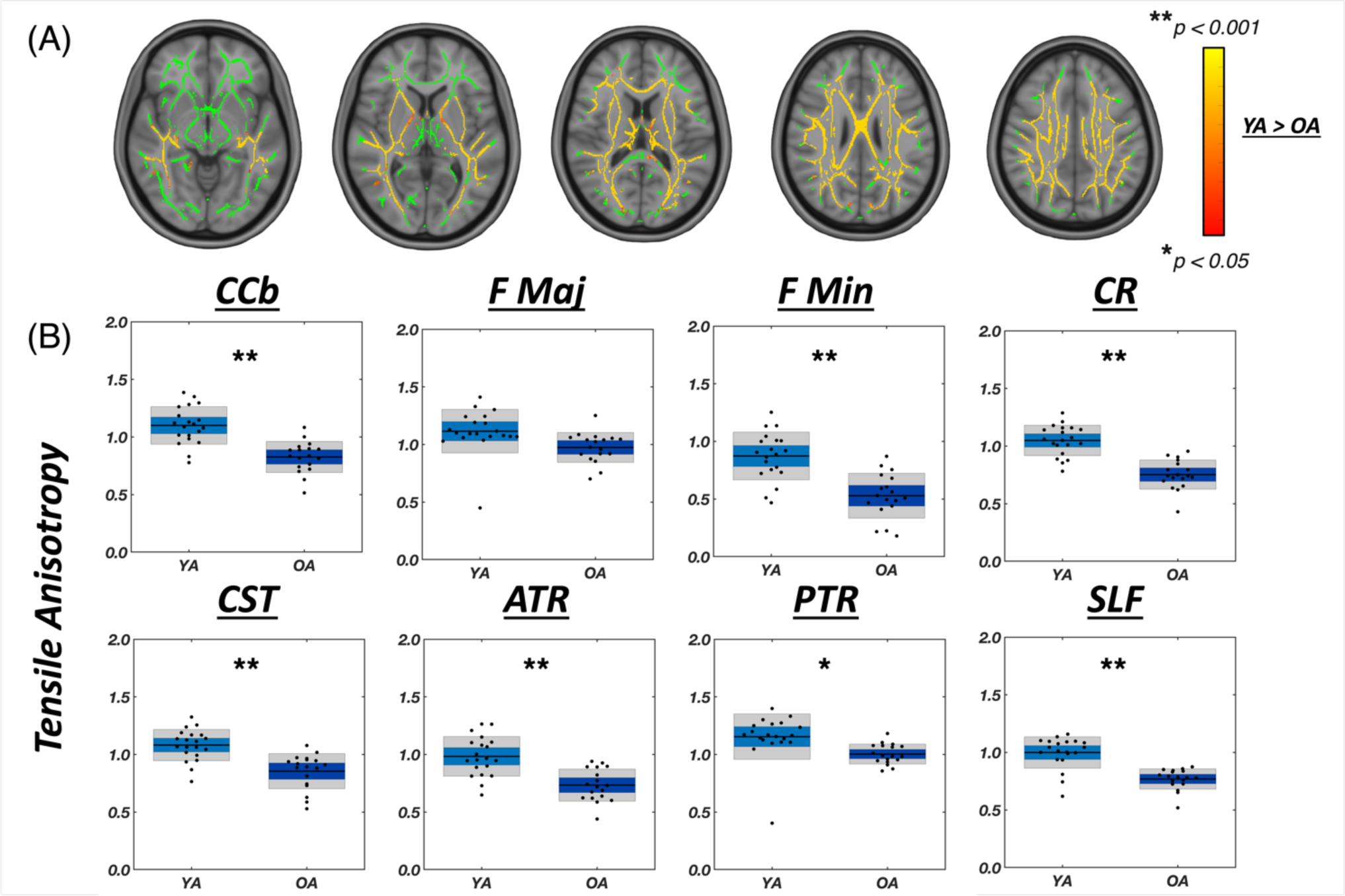
(A) Results from voxel-wise statistical analysis comparing younger adults (YA) versus older adults (OA) for substrate shear stiffness (ζ), depicted for several axial slices (Z = -6, 5, 14, 26, 35). Significant voxels from the YA>OA contrast are highlighted in red-yellow, overlayed on the white matter skeleton shown in green. (B) Segmented white matter region averages of tensile anisotropy, with asterisks indicating level of statistical significance for two-sample student t-tests with Bonferroni correction, * p<0.00625, ** p<0.001.

### 3.4 Contrasting MRE and DTI parameters using TBSS

In addition to evaluating the individual MRE parameters, we contrasted the anisotropic MRE measures with diffusion tensor imaging (DTI) parameters to investigate potential overlap and differences in their sensitivity to age-related changes in WM (Figure 6). The percentage of significant voxels exhibiting age-related differences (Younger Adult > Older Adult; p < 0.05) was calculated for each parameter across the white matter skeleton of the cerebrum. Tensile anisotropy (ζ) exhibited the highest percentage of significant voxels (59.8%), followed by fractional anisotropy (FA) from DTI (59.7%), shear anisotropy (ϕ) (41.8%), and substrate shear stiffness (μ) (6.1%). The overlap between significant voxels in FA and ϕ was 27.6%, while the overlap between FA and ζ was 39.5%. These moderate levels of overlap suggest that the age-related effects on anisotropic MRE parameters capture distinct aspects of microstructural changes compared to DTI-derived measures of diffusion anisotropy.

**Figure 6:**
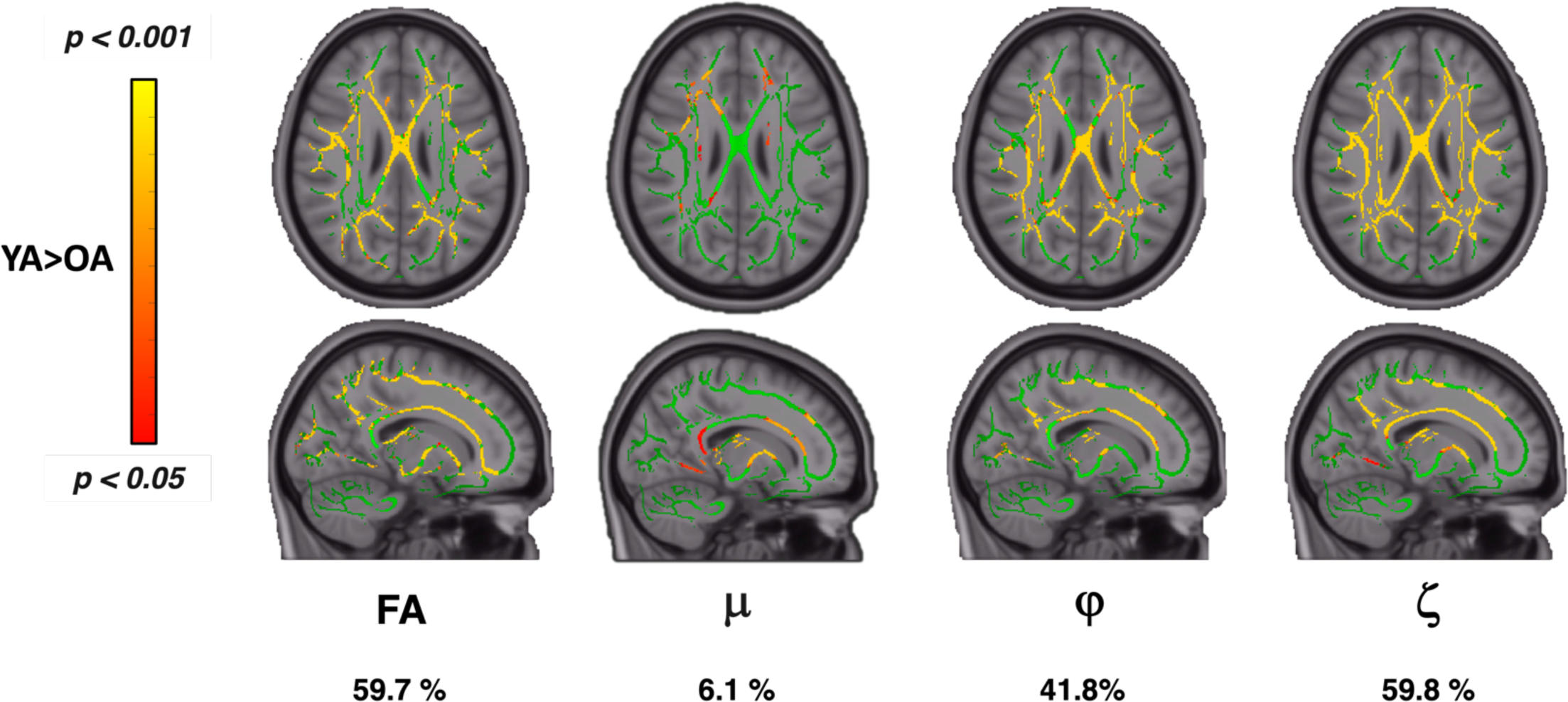
Summary of TBSS voxel-wise statistical analysis for fractional anisotropy and all mechanical parameters comparing young and older adults. Shear stiffness has the smallest percentage of significant voxels within the cerebrum white matter, whereas both anisotropy parameters have comparable values relative to fractional anisotropy.

To assess the added value of MRE anisotropy parameters beyond DTI-derived FA, we further performed logistic regression analyses using ROI-derived values. As shown in Table 2, both shear anisotropy and tensile anisotropy exhibited significant effects, indicating that these MRE measures indeed provide complementary information to FA in discriminating between younger and older adults. Overall, these findings suggest that anisotropic MRE parameters, particularly tensile anisotropy, are sensitive to age-related changes in WM microstructure and offer unique insights beyond conventional DTI measures.

**Table 2:** Summary of logistic regression outputs including p-values and Beta coefficients for analysis performed in each white matter tract. Significance level of p < 0.05 is denoted by the bolded values.

| *Region* | **FA, $\mu$** |  | **FA, $\phi$** |  | **FA, $\zeta$** |  |
| --- | --- | --- | --- | --- | --- | --- |
|  | p-value | Beta | p-value | Beta | p-value | Beta |
| **CCB** | 0.938 | 0.000 | **0.019** | 12.476 | **0.016** | 23.798 |
| **Fmaj** | 0.341 | 0.002 | 0.076 | 7.272 | **0.061** | 5.408 |
| **Fmin** | **0.015** | 0.007 | **0.011** | 34.596 | **0.022** | 15.379 |
| **CR** | 0.179 | 0.004 | **0.004** | 73.000 | **0.004** | 19.679 |
| **CST** | 0.539 | 0.001 | **0.004** | 22.102 | **0.014** | 28.675 |
| **ATR** | **0.021** | 0.006 | **0.010** | 84.193 | **0.014** | 18.213 |
| **PTR** | 0.115 | 0.003 | 0.265 | 4.859 | **0.032** | 7.637 |
| **SLF** | 0.431 | 0.002 | **0.003** | 51.069 | **0.003** | 18.881 |

## 4. DISCUSSION

This paper presents a cross-sectional study examining mechanical properties of WM tracts in the aging brain through application of anisotropic MRE with TI-NLI using wave fields acquired via multi-excitation MRE. Our investigation focuses on the substrate shear stiffness, shear anisotropy, and tensile anisotropy of WM in older adults in comparison with younger adults. Analysis of mechanical properties in the two groups revealed significant differences in several white matter regions, with the older group exhibiting slightly lower stiffness than the younger group but much lower mechanical anisotropy, likely reflecting lower WM integrity in older adults. Notably, our results reveal minimal overlap between MRE measures of mechanical anisotropy with fractional anisotropy from diffusion MRI, providing evidence that these measures are independent and convey distinct information in characterizing WM integrity in aging.

Despite the commonly reported lower tissue stiffness in age, we observed only a modest 5.0% average lower stiffness μ in older adults compared to their younger counterparts, with equivalent annual differences between -0.002 and -0.008 kPa/year. Only a small number of voxels from TBSS exhibited significantly lower substrate shear stiffness in older adults, and only the forceps minor and anterior thalamic radiation ROIs had statistically significantly lower stiffness. This lower stiffness appears less pronounced than previous studies employing isotropic material models in global or gray matter regions, with reported decreases in cerebral stiffness ranging from -0.008 to -0.025 kPa/year (Arani et al., 2015; Lv et al., 2020; Sack et al., 2009; Takamura et al., 2020). Using isotropic nonlinear inversion methods, Delgorio et al. (2021) reported an annual decrease in hippocampal stiffness of -0.014 kPa/year, while Hiscox et al. (2018) reported -0.006 kPa/year, with up to 24% difference between younger and older groups in specific subcortical regions. Compared to stiffness, age-related effects on mechanical anisotropy were much greater. Shear anisotropy (ϕ) was lower in older adults by 32.5% on average, and tensile anisotropy (ζ) was lower by 24.48%, with nearly all tracts exhibiting significant anisotropy differences between groups, showcasing a higher sensitivity to age-related WM changes than shear stiffness.

The diminished anisotropy likely explains the discrepancy between only moderately lower stiffness in our study compared to larger aging effects in previous studies employing isotropic models. Considered differently, stiffness *perpendicular* to the fibers was not strongly affected by age, but stiffness *parallel* to the fibers was significantly lower in the older group, in general, and becomes more similar to perpendicular stiffness. Thus, we can interpret our findings as largely driven by axonal factors, as we would expect fiber characteristics to affect stiffness most strongly in the direction of the fibers. Isotropic inversions of anisotropic tissues generally return some composite average of direction-dependent properties depending on propagation and polarization directions of the waves in the displacement fields (Anderson et al., 2016; Smith et al., 2020; Tweten et al., 2017). Hence, our reported stiffness and age-related effects on stiffness are both lower with anisotropic MRE compared to isotropic MRE where tissue may appear stiffer depending on wave characteristics.

Our findings align with an extensive literature investigating age-related changes in WM microstructure through diffusion MRI, where lower fractional anisotropy indicative of reduced WM integrity is observed in older adults (Pfefferbaum et al., 2000). Subsequent research consistently revealed regional vulnerabilities, with an anterior-posterior gradient of susceptibility (Davis et al., 2009; Head et al., 2004) and greater differences in WM tracts which connect to the prefrontal cortex (Michielse et al., 2010), consistent with the “last in, first out” hypothesis, wherein anterior tracts that mature latest during development show the earliest age-related decline (Davis et al., 2009; Lebel et al., 2012). While our data suggests lower mechanical properties with age throughout WM, especially tensile anisotropy, anterior WM appears to be most strongly or consistently affected, i.e. from results of the TBSS analysis of stiffness. We note that the anterior thalamic radiation and forceps minor – both anterior tracts – were the only regions we examined to exhibit significantly lower stiffness, shear anisotropy, and tensile anisotropy in the older group.

While our study did not directly assess cognitive performance, our results suggest potential implications for cognitive function based on previous associations between white matter integrity and cognition. The observed alterations in mechanical properties in regions such as the body of the corpus callosum and forceps minor may have implications for interhemispheric communication and cognitive processes that rely on efficient information transfer between hemispheres, such as attention, executive function, and memory (Pfefferbaum et al., 2000). Furthermore, reductions in tensile anisotropy observed in the anterior thalamic radiation and corticospinal tract align with regions previously implicated in cognitive function decline, particularly in domains such as processing speed, attention, and motor function (Bennett & Madden, 2014; Madden et al., 2009). By highlighting these regional associations, our study adds to the growing body of evidence emphasizing the importance of considering regional variations in WM integrity and their potential impact on cognitive aging processes. Furthermore, our results underscore the value of combining mechanical property assessments with traditional diffusion metrics to gain a more comprehensive understanding of the underlying structural and functional changes contributing to cognitive decline.

One previous study used an anisotropic MRE scheme to study brain stiffness during aging. Kalra et al. (2019) analyzed subjects aged 18-62 years using a 60 Hz MRE acquisition with 2.5 mm voxel resolution. However, they did not find significant correlations with age for anisotropic parameters in any of their WM regions of interest, and instead only showed lower stiffness with age in gray matter but without a change in anisotropy. Their anisotropic MRE approach is based on direct inversion with a large number of anisotropic parameters, which may induce uncertainty in estimated outcome measures. Conversely, our TI-NLI approach can effectively describe both the heterogeneity in properties and fiber direction and recovers only a minimal number of parameters, as described in previous studies (Jyoti et al., 2022; McGarry et al., 2022). Additionally, the use of TI-NLI with multi-excitation MRE has been well-characterized and shown to yield repeatable results in WM tracts (Smith et al., 2022). This robust methodology, and improved resolution, enhances the reliability of our findings and provides a more comprehensive understanding of the changes in anisotropic mechanical properties of WM tracts with age.

Wang et al. (2023) also considered age-related effects on anisotropic MRE parameters, but instead examined minipig brains across a period of development. That study also used TI-NLI and an actuator modified from Guertler et al. (2018) to achieve multiple wave fields as in human multi-excitation MRE. They observed higher shear stiffness, shear anisotropy, and tensile anisotropy in WM compared to gray matter. While they found a small, statistically significant increase in substrate shear stiffness with age, no significant effects of age were evident on shear or tensile anisotropy. Notably, tensile anisotropy exhibited a slight, non-significant decreasing trend with age in WM, while anisotropy might be expected to increase with development due to rapid myelination during this time period. The authors speculate that this unexpected trend may be related to the effects of myelination and cytoskeletal reorganization on the mechanical anisotropy. Compared to consistent results regarding brain stiffness in older age, mechanical properties during brain development and maturation are complex and incompletely understood (McIlvain et al., 2018; McIlvain, Schneider, et al., 2022; Ozkaya et al., 2021; Yeung et al., 2019).

A common question in the field of brain MRE is whether mechanical property estimates are unique from other microstructural imaging metrics such as those from diffusion MRI, especially in WM where FA is commonly used as a measure of WM integrity. Our data suggest that MRE provides unique, complementary information, as evidenced by our logistic regression results. Even though both MRE and diffusion MRI are sensitive to WM microstructure, and both are affected by aging processes (Davis et al., 2009), we can interpret these findings as arising from differences in sensitivity to varying aspects of the underlying microstructure between the methods. For example, anisotropic MRE may better capture or resolve aspects of the extra-axonal matrix than FA. These results also provide evidence that mechanical anisotropy estimates from TI-NLI are not strongly influenced by the underlying data from diffusion used during inversion. This finding is consistent with previous work by McGarry et al. (2022) that demonstrated that adding noise to input eigenvector directions did not overly corrupt TI-NLI property estimates. Hence, the mechanical anisotropy parameters may offer supplementary information on WM microstructure beyond what diffusion MRI alone can provide.

A limitation of the present study is the cross-sectional design and modest sample size (20 younger adults, 18 older adults). Longitudinal, repeated anisotropic MRE measurements across the lifespan in a larger cohort would be necessary to fully characterize the trajectories of WM mechanical anisotropy changes with aging. We compared the anisotropic MRE parameters with just FA from DTI, but many more advanced diffusion models are available that may provide sensitivity to aspects of tissue microstructure that are captured by MRE. Future studies on WM tracts should examine relationships between MRE and more advanced diffusion models, such as multi-shell or crossing fiber models and diffusion kurtosis imaging. Furthermore, how mechanical anisotropy reflects the underlying neurobiological mechanisms related to declining WM integrity in aging, such as demyelination, oligodendrocyte death, or axonal death, remain unclear and animal studies are needed that can study mechanical and cellular metrics simultaneously to understand the sensitivity of MRE outcomes.

## 5. CONCLUSION

This study demonstrates the capability of anisotropic MRE to characterize age-related differences in the mechanical properties of WM tracts in the human brain. These age-related alterations in anisotropic mechanical properties likely reflect underlying microstructural changes associated with aging, such as demyelination, axonal degeneration, and extracellular matrix changes. Importantly, the observed differences in mechanical anisotropy provide complementary information to traditional diffusion MRI measures, suggesting that anisotropic MRE offers a novel perspective on WM integrity. By combining anisotropic MRE with diffusion MRI and other neuroimaging techniques, we can gain a more comprehensive understanding of the structural and functional changes that occur in the aging brain. Ultimately, this approach may lead to improved diagnostic and prognostic capabilities for age-related neurodegenerative conditions and other neurological disorders affecting WM in the elderly.

## 6. ACKNOWLEDGEMENTS

Funding for this work was provided, in part, by grants from the National Institutes of Health (R01-EB027577, R01-AG05883, and U01-NS112120) and the Office of Naval Research (N00014-22-1-2198).

